# The genome of the CTG(Ser1) yeast S*cheffersomyces stipitis* is plastic

**DOI:** 10.1101/2021.02.15.431239

**Authors:** Samuel Vega Estevez, Andrew Armitage, Helen J. Bates, Richard J. Harrison, Alessia Buscaino

**Affiliations:** University of Kent, School of Biosciences, Kent Fungal Group, Canterbury Kent, CT2 7NJ, UK; Natural Resources Institute, University of Greenwich, Chatham Maritime, Kent, ME4 4TB, UK; NIAB Cambridge Crop Research, 93 Lawrence Weaver Road, Cambridge CB3 0LE

## Abstract

Microorganisms need to adapt to environmental changes, and genome plasticity can lead to rapid adaptation to hostile environments by increasing genetic diversity. Here, we investigate genome plasticity in the CTG(Ser1) yeast *Scheffersomyces stipitis*, an organism with an enormous potential for second-generation biofuel production. We demonstrate that *S. stipitis* has an intrinsically plastic genome and that different *S. stipitis* isolates have genomes with distinct chromosome organisation. Real-time evolution experiments show that *S. stipitis* genome plasticity is common and rapid as extensive genomic changes with fitness benefits are detected following *in vitro* evolution experiments. Hybrid MinION Nanopore and Illumina genome sequencing identifies retrotransposons as major drivers of genome diversity. Indeed, the number and position of retrotransposons is different in different *S. stipitis* isolates, and retrotransposon-rich regions of the genome are sites of chromosome rearrangements. Our findings provide important insights into the adaptation strategies of the CTG (Ser1) yeast clade and have critical implications in the development of second-generation biofuels. These data highlight that genome plasticity is an essential factor to be considered for the development of sustainable *S. stipitis* platforms for second-generation biofuels production.

## INTRODUCTION

Eukaryotic genomes are often described as stable structures with well-preserved chromosome organisation, and genome instability is viewed as harmful. However, an increasing body of evidence demonstrates that eukaryotic microorganisms have a plastic genome and genome instability is instrumental for rapid and reversible adaptation to hostile environments (1–4). This is because genomic instability can increase genetic diversity, allowing the selection of genotype(s) better adapted to a new environment (5, 6). Repetitive DNA elements are major contributors to genome plasticity as repeats can undergo inter and intra-locus recombination, resulting in gene conversion, gross chromosomal rearrangements and segmental aneuploidies (7). Transposable Elements (TE), a specific class of repetitive elements, alter genome organisation by recombination-dependent mechanisms and by jumping to new sites in the genome (8). TEs belong to two major classes: DNA transposons (Class II) and retrotransposons (Class I). DNA transposons utilise a “cut and paste” mechanism in which the parental element excises from its original location before integrating elsewhere (9). In contrast, retrotransposons replicate through reverse transcription of their RNA and integrate the resulting cDNA into another locus. Retrotransposons can be further classified into Long Terminal Repeats (LTR) retrotransposon and non-LTR retrotransposons (10). LTR retrotransposons are characterised by two LTR sequences flanking an internal coding region containing the genes encoding for the structural protein GAG and enzyme POL required for reverse transcription and integration (11). While POL enzymes are conserved across organisms, GAG proteins are poorly conserved (12). LINE elements are one of the most abundant non-LTR retrotransposons and they are typically composed of a 5’ non-coding region, two ORFs (ORF1 and ORF2) and a 3’ non-coding region that is marked by a poly(A) tail (13). ORF1 proteins have a diverse amino acid sequence, but they often contain a DNA-binding motif (14). ORF2 encodes endonuclease and reverse transcriptase activity that are critical for transposition (15).

The CTG (Ser1) clade of fungi, in which the CTG codon is translated as serine rather than leucine, is an important group of ascomycetous yeasts featuring yeasts that hold great promises in biotechnology, such as *Scheffersomyces stipitis*, and dangerous human fungal pathogens, such as *Candida albicans* (16). The CTG(Ser1) clade comprises several species with different lifestyles and genomic organisations, including haploid and diploid species that colonise diverse environments by reproducing sexually or para-sexually (16–19). One common feature of CTG(Ser1) species is their ability to adapt remarkably well to extreme environments (20). For example, CTG(Ser1) yeasts can grow on various carbon sources and are highly tolerant to environmental changes such as changes in osmolarity (16, 19, 20). It is well established that genome plasticity is a critical adaptive mechanism in the human fungal pathogens *Candida albicans*, the most studied CTG (Ser1)-clade member (4). In *C. albicans*, stress increases genome instability by affecting the rate and type of genomic rearrangements (21). Different classes of DNA repeats drive this genetic variation, including TEs, long repeats and Major Repeat Sequences (MRS) (22–24). It is still unknown whether genome plasticity is a general feature of the CTG(Ser1) clade and whether DNA repeats are drivers for genome diversity across this yeast group.

This study investigates genome plasticity in *S. stipitis*, a CTG (Ser1)-clade yeast with great potential for the eco-friendly and ethical production of second-generation biofuels (25–27). Second-generation biofuels are generated by fermentation of lignocellulose biomass, produced in large amounts (>1.3 billion tons produced annually) as waste following agricultural and forestry processing operation (27). Lignocellulose is a heteropolymer composed of fermentable hexose sugars, such a glucose, and pentose sugars, such as xylose (28). The yeast *Saccharomyces cerevisiae*, usually the organism of choice for industrial production of ethanol, is not suitable for the production of second-generation ethanol because it cannot ferment pentose sugars as it lacks specific transporters and enzymatic network important for their metabolism (28). *S. stipitis* holds excellent potential for biofuel derived from green waste because it is one of the few yeast species that can ferment both hexose and pentose sugars (25–27). *S. stipitis* is a non-pathogenic haploid yeast that is found in the gut of wood-ingesting beetles, in hardwood forests or areas high in agricultural waste (29). Contrary to *C. albicans*, *S. stipitis* has a canonical sexual cycle whereby mating of haploid cells generate diploid cells that undergo meiosis and produce haploid spores (30). Although several *S. stipitis* natural isolates are used for the optimisation of second-generation biofuels production, the genome of only one strain (Y-11545) has been sequenced and assembled to the chromosomal level (31). The Y-11545 genome has a size of 15.4 Million base pair (Mbp) organised in 8 chromosomes and containing ~6000 protein-coding genes (31–33). *S. stipitis* chromosomes are marked by regional centromeres composed of full-lenghts LTR retrotrasposons (Tps5a, Tps5b and Tps5c) and non-coding, non-autonomous LARD (large retrotransposon derivative) elements (31, 33).

To investigate the plasticity of the *S. stipitis* genome, we have taken several complementary approaches. Firstly, we systematically identified *S. stipitis* DNA repeats and investigated the genotypic diversity of 27 different *S. stipitis* natural isolates collected from different environments. Secondly, we combined MinION Nanopore with Illumina genome sequencing to generate a high-quality chromosome-level sequence assembly of a second *S. stipitis* natural isolate (Y-7124) and compared its genome structure to the reference Y-11545 genome. Lastly, we performed *in vitro* evolution experiments and analysed *S. stipitis* genome organisation changes following laboratory passaging under stress or unstressed growth conditions. Thanks to this combined approach, we discovered that the *S. stipitis* genome is plastic. Genome plasticity is not regulated by stress, however large chromosome rearrangements are linked to adaptation to hostile environments. We demonstrate that different *S. stipitis* natural isolates have distinct chromosomal organisations and that transposable elements drive this extensive intra-species genetic variation. Our findings have important implications for second-generation biofuel production as genome plasticity is a paramount factor to be considered for the successful development of superior biofuel-producer *S. stipitis* strains.

## MATERIAL AND METHODS

### Yeast strains and Growth Conditions

Strains were obtained from the Agricultural Research Service (ARS) Collection, run by the Northern Regional Research Laboratory (NRRL) (Peoria, Illinois, USA), or the National Collection of Yeast Cultures (NCYC) (Norwich, United Kingdom) (**Table S1)** and confirmed by sequencing (primers AB798 and AB799 of the 26S rDNA (D1/D2 domain) (34) (**Table S2**). Routine culturing was performed at 30 °C with 200 rpm agitation on Yeast Extract-Peptone-D-Glucose (YPD) media. Phenotypic and *in vitro* evolution analyses were conducted on Synthetic Complete (SC) media containing glucose (SC-G), xylose (SC-X), or a mixture of 60% glucose and 40% xylose (SC-G+X). SC-G was used as a reference media as glucose is the preferred carbon source for both the model system *S. cerevisiae* and *S. stipitis*, SC-X was used because of *S. stipitis* unique ability to utilise xylose as a carbon source and SC G+X was used because this sugar combination resemble the ratio found in lignocellulose (28). Uridine (0.08 g/L in YPD and SC) and adenine hemisulfate (0.05 g/L in YPD) were added as growth supplements. Solid media were prepared by adding 2% agar.

### Contour-clamped homogeneous electric field (CHEF) electrophoresis

Intact yeast chromosomal DNA was prepared as previously described (35). Briefly, cells were grown overnight and spheroplast were prepared in an agarose plug by treating cells (~ OD_600_=7) with 0.6 mg/ml Zymolyase 100T (Amsbio #120493-1) in 1% Low Melt agarose (Biorad® # 1613112). Chromosomes were separated in a 1% Megabase agarose gel (Bio-Rad) in 0.5X TBE using a CHEF DRII apparatus. Run conditions as follows: 60-120s switch at 6 V/cm for 12 hours followed by a 120-300s switch at 4.5 V/cm for 12 hours, 14 °C. Chromosomes were visualised by staining the gel 0.5x TBE with ethidium bromide (0.5 μg/ml) for 30 minute, followed by destaining in water for 30 minutes. Images were capture using a Syngene GBox Chemi XX6 gel imaging system.

### Southern Blotting

DNA from CHEF gels were transferred overnight to a Zeta-Probe GT Membrane (Biorad®, #162-0196) in 20x SSC and crosslinked using UV (150 mJ). Probing and detection of the DNA were conducted as previously described (36). Briefly, probes were generated by PCR incorporation of DIG-11-dUTP into target sequences following manufacturer’s instructions (Roche). Chromosome 5-chromosome 7 translocation was detected using primers AB1028 and AB1029 amplifying a 180 bp region of chromosome 5 (Chr5 nt: 448,855-449,034) in Y-11545 and in chromosome 7 of Y-7124 (Chr7 nt: 494,698-494,877) (**Table S2**). The membrane was hybridised overnight at 42 °C with DIG Easy Hyb (Roche®, 11603558001). The DNA was detected with anti-digoxigenin-Alkaline Phosphatase antibody (Roche®, #11093274910) and CDP Star ready to use (Roche®, #12041677001) according to manufacturer instructions.

### Phenotypic characterisation

Growth analyses were performed using a plate reader (SpectrostarNano, BMG labtech) in 96 well plate format at 30 °C for 48 hours in SC-G, SC-X or SC-G+X. The growth rate (μ, hours^−1^) was calculated using: μ = (ln(X_2_)-ln(X_1_))/(t_2_-t_1_), where: (i) X_1_ is the biomass concentration (OD_600_) at time point one (t_1_, lag time) (ii) X_2_ is the biomass concentration (OD_600_) at time point two (t_2_, end of exponential growth phase). The maximum OD (OD units) was determined with the MAX() from Excel (Microsoft®). The lag time (minutes) was determined visually as the time in which the exponential growth starts. Experiments were performed in 3 technical and 3 biological replicates. Heatmaps showing the average of 3 biological replicates were generated by R using the library *pheatmap*. ANOVA test was performed to study differences on growth rate, maximum OD and lag time between the strains. The equality of variances presumption was tested using Levene’s test, whereas the normality of the data was tested by Shapiro-Wilk. When both assumptions were satisfied, a Tukey’s honest significant test was used to determine significant differences between the natural isolates and the reference Y-11545 strain. When assumption of equal variance were violated, one-way test was used to indicate significance. In the case of equal variances, but a non-normal distribution of data, the Kruskal-Wallis rank sum test was used to indicate statistical differences and significance was determined by pairwise testing. A p-value lower than 0.05 was considered significant for all these statistical tests.

### Adaptive Laboratory Evolution

A single colony of the *S. stipitis* strain NRRL Y-7124 was grown overnight in 5 ml of YPD at 30 °C, plated in YPD at a cell density of 100 and grown 48 hours at 30 °C. 36 single colonies were streaked in two SC-G+X plates and grown at 30 °C and 37 °C, respectively and streaked daily for a total of 56 passages (8 weeks). The karyotype variability of the colonies was assessed by CHEF electrophoresis. Phenotypic differences were assessed by spotting assays. Strains with rearrangements were grown overnight is SC-G+X and were diluted to an OD_600_=1. From this, five 1/10 serial dilutions were prepared and the cells were plated in SC-G+X using a replica plater (Sigma Aldrich, R2383-1EA) and grown for 48 hours at both 30 °C and 37 °C. Strains with no karyotypic modifications after evolution were also used as control.

### Identification of DNA repeats

Long sequences (>100 nucleotides) present more than once in the Y-11545 and Y-7124 genomes were identified by aligning each genome to itself using BLASTN. Repetitive elements (E < 1e-04) were manually verified using IGV/SNAPGene, and clustered repeats were combined. This repeats dataset was manually examined to further classify it as (a) related to transposable elements (b) telomeric repeats, (c) centromeres (d) belonging to protein coding gene families and (e) MRS repeats. Transposons were classified using established guidelines (10). Briefly, LTR-transposons were identified by detecting two Long-terminal Repeat sequences (size 260-430 nt) flanking an internal coding region. These potential LTR-transposons were further annotated for the presence of the following marks: LTR flanked by a TG and CA di-nucleotides, presence of a Primer Binding Site (PBS) with homology to *S. stipitis* tRNAs (GtRNAdb (http://gtrnadb.ucsc.edu/index.html), presence of a coding region with homology to *pol* gene and containing an Integrase (INT), Reverse Transcriptase (RT) and RNAse H (RH) domain. Non-LTR LINE transposons were identified by detection of coding regions homologous to LINE retrotransposons ORF1 (containing a Zn-finger), ORF2 (containing an Endonuclease and a Reverse Transcriptase domain) and terminal Poly-A sequence.

Retrotransposons were classified into different families based on sequence similarity with a 90% cut-off. Terminal telomeric tandem repeats were identified using Tandem Repeats Finder (37) with default parameters. Regional centromeres were identified based on them being the only regions of the genome with a large retrotransposon Tps5 cluster (~ 20-40 kb) as previously described (33). Gene families were identified by extracting coding-regions from our repeats datasets and performing Clustal Omega sequence alignment and PFAMs domain identification using SMART(http://smart.embl.de) (38). The identified gene families were compared to published information (39). The presence of MRS repeats was explored using BLASTN and by searching for clusters of non-coding tandem repeats, a hallmark of *C. albicans* MRS, with no-homology to retrotransposons and not-located at chromosome ends. Sequence alignments were visualised with Jalview v2.11.1.0 (40). Phylogenetic trees were generated with phyloT : a phylogenetic tree generator (biobyte.de) using default parameters and visualised with Itol (https://itol.embl.de/).

### Genome sequencing

The genome of *S. stipitis* isolate Y-7124 was sequenced by Illumina short-read and MinION long-read technologies. To this end, DNA was extracted from an overnight culture using the QIAGEN genomic tip 100/G kit (Qiagen®, #10243) according to manufacturing protocol. For long-read sequencing, MinION (Oxford Nanopore, Oxford UK) was performed on a DNA library prepared from size selected gDNA. DNA fragments greater than 30 Kb were selected using a Blue Pippin (Sage Science) and concentrated using Ampure beads. From this, a DNA library was prepared using a Ligation Sequencing Kit 1D (SQK-LSK108) and run on the Oxford Nanopore MinION flowcell FLOMIN 106D. The same gDNA extract was also used for the preparation of Illumina libraries. In this case, the DNA was sheared using the Covaris M220 with microTUBE-50 (Covaris 520166) and size selected using the Blue Pippin (Sage Science). The library was constructed using a PCR-free kit with NEBNext End Repair (E6050S), NEBNext dA-tailing (E6053S) and Blunt T/A ligase (M0367S) New England Biolabs modules. Sequencing was performed on a MiSeq Benchtop Analyzer (Illumina) using a 2×300bp PE (MS-102-3003) flow cell.

### Genome assembly

Base-calling and demultiplexing were conducted with Albacore v2.3.3 (available at https://community.nanoporetech.com). Adapters and low-quality data were trimmed using the eautils package fastq-mcf 1.04.636 (https://expressionanalysis.github.io/ea-utils/). On nanopore sequence data, adapter trimming was performed with Porechop v.0.1.0 (https://github.com/rrwick/Porechop). Genome assembly was completed using long reads, with read correction performed with Canu v1.8 (41) followed by assembly in SMARTdenovo github commit id 61cf13d (42)). The draft assembly was corrected using the corrected nanopore reads through five rounds of Racon github commit 24e30a9 (43), and then by raw fast5 files using 10 rounds of Nanopolish v0.9.0 (44). Illumina sequencing reads were then used to polish the resulting assembly through 10 rounds of Pilon v1.17 (45). Following genome assembly, BUSCO v3 was run to assess evolutionary conserved gene content (46), using the Saccharomycetales_odb9 gene database. The Saccharomycetales database contains 1711 genes, which are therefore expected to be present in *S. stipitis*. Of these, 1683 (98.36%) were identified in the Y-7124 assembly demonstrating a good level of completeness (>95%) (**Table S8**). Assembly size and contiguity statistics were assessed using QUAST v4.5 (47). This initial assembly of the nuclear genome contained 10 contigs. A chromosome level assembly was produced by identification of overlapping regions between the contigs: a 244 Kbp overlapping region between contig 7 and 2 led to the final assembly of Chromosome 1, a 83 Kpb overlapping region between contig 9 and 10 led to the final assembly of Chromosome 8.

### Genome annotation

Genome annotation was performed using FUNGAP v1.0.1 (48) with fastq reads from NCBI SRA accession SRR8420582 used as RNA-Seq training data and protein sequences taken from NCBI assembly accession GCA_000209165.1 for *S. stipitis* NRRL Y-11545 (CBS6054) used for example proteins. Protein fasta files were extracted from predicted gene models using the yeast mitochondrial code (code 3) and the alternative yeast nuclear code (code 12). Functional annotation of gene models was performed through BLASTp searches vs all proteins from the NCBI reference fungal genomes (downloaded 11^th^ April 2020), retrieving the top-scoring blast hit with an E-value < 1×10^−30^. These annotations were supplemented with domain annotations from Interproscan v5.42-78.0 (49). The annotated genome was submitted to NCBI, with submission files prepared using GAG v2.0.1 (http://genomeannotation.github.io/GAG.), Annie github commit 4bb3980 (http://genomeannotation.github.io/annie) and table2asn_GFF v1.23.377 (available from https://ftp.ncbi.nih.gov/toolbox/ncbi_tools/converters/by_program/tbl2asn/).

### Comparative genomics

Whole-genome alignment between Y-7124 and Y-11545 was performed using the nucmer tool from the MUMmer package v4.0 (50) with results visualised using Circos v0.6 (51). Orthology analysis was performed between predicted proteins from these isolates using OrthoFinder v2.3.11 (52), with results visualised using the package VennDiagram in R (53).

Sequence variants were identified in Y-7124 through comparison to the Y-11545 assembly. Short read sequence data for Y-7124 were aligned to the reference genome using BWA v 0.7.15-r1140 (54), before filtering using using picardtools v2.5.0 to remove optical duplicates (http://broadinstitute.github.io/picard/). SNP and insertion/deletion (InDel) calling was performed using GATK4 (55). Low confidence variants were then filtered using VCFtools v0.1.15 (56) using minimum mapping quality of 40, phred quality of 30, read depth of 10 and genotype quality of 30. Effect of variants on NRRL Y-11545 gene models was determined using SnpEff v4.2 (57).

## RESULTS

### Classification of *S. stipitis* DNA repeats

DNA repeats are drivers of genome variation. Understanding the repertoire of repetitive elements associated with a genome is critical to gain insights into the genome diversity of a specific organism. Comparative genomic analyses have identified different repetitive elements in some CTG(Ser1) clade members, yet a comprehensive survey of *S. stipitis* repetitive elements is lacking (18, 58). Therefore, we sought to classify the major classes of repetitive elements associated with the Y-11545 sequenced genome by aligning the genomic sequence of each strain to itself and identifying long sequences (>100 nucleotides) present more than once in the genome. The genomic position of these repeats was manually verified, and clustered repeats were combined and categorised depending on their genomic position, structure and sequence similarity. Our analyses identified known *S. stipitis* repeat-rich loci such as centromeric transposon-clusters, the NUPAV sequence, an integrated L-A ds-RNA virus, and several gene families (32, 33, 59). As observed in other members of the CTG (Ser1)-clade, we did not detect any MRS repeats, a class of repetitive elements found only in *C. albicans* and the closely related *C. dubliensis* and *C. tropicalis* species (58, 60). Here we focus on intra-chromosomal or inter-chromosomal repeats that have not been described to date: non-centromeric TEs, subtelomeric regions and telomeric repeats (**Fig 1**).

**Figure 1.**
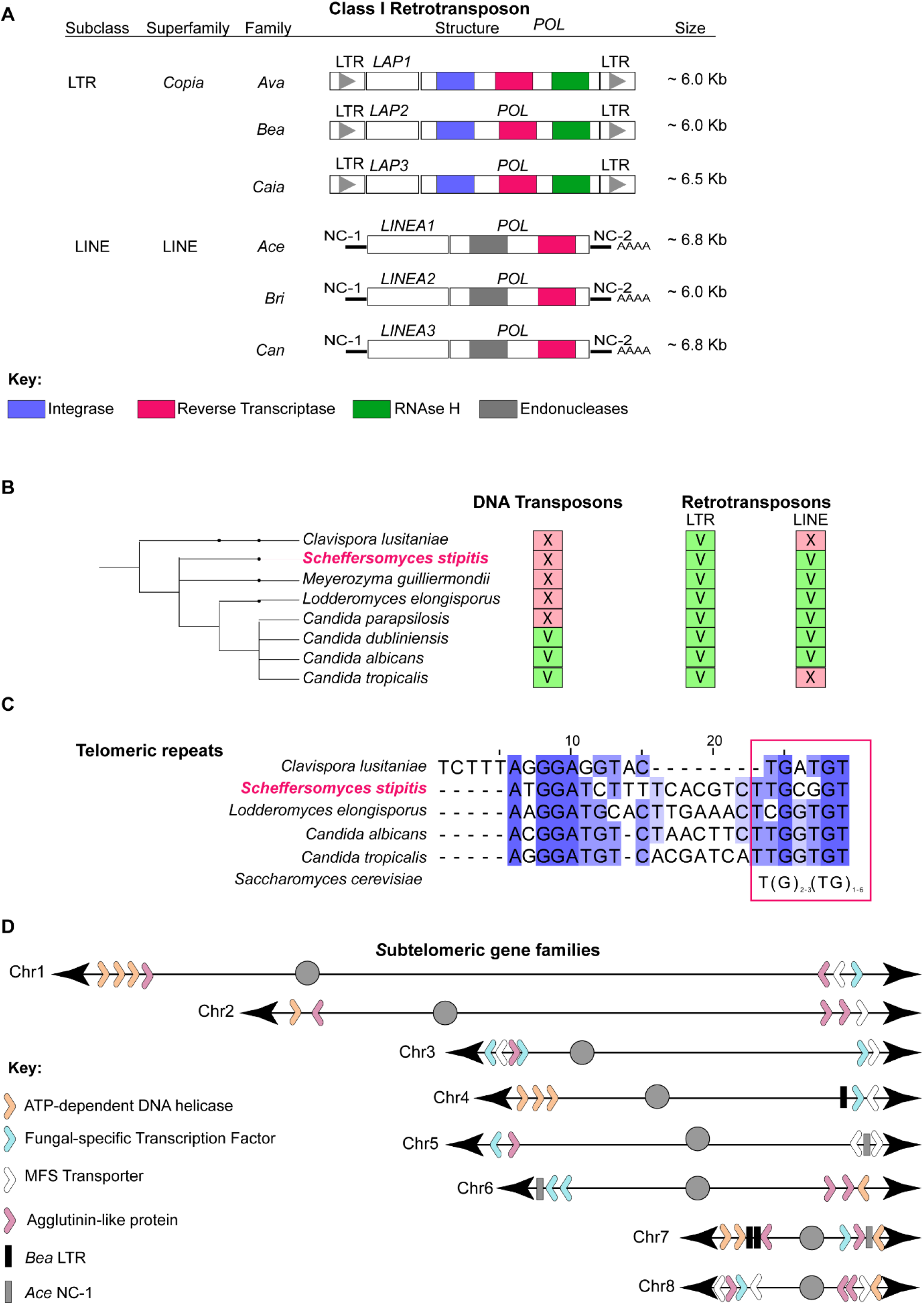
Classification of non-centromeric *S. stipitis* repeats **A)** Schematics of non-centromeric retrotransposons identified in this study. For each transposon, subclass, superfamily and family is indicated. The organisation of coding and non-coding sequences of each transposon is displayed. **B)** Cladogram showing CTG (Ser1)-clade species with known transposable elements (this study and (60, 68). The presence (V) or absence (X) of a TE is indicated. **C)** Sequence alignment of telomeric terminal repeats in members of the CTG (Ser1)-clade (*C. lusitaniae*, *S. stipitis*, *L. elongisporus*, *C. albicans* and *C. tropicalis*) (this study and (60, 68). Consensus sequence to the *S. cerevisiae* telomeric repeats is indicated (Magenta box). **D)** Schematics of gene family members associated with *S. stipitis* subtelomeres (30 Kb from chromosome end).

We identified six novel retrotransposon families scattered along chromosome arms: 3 LTR retrotransposons (*Ava*, *Bea* and *Caia*) and 3 LINE retrotransposons (*Ace*, *Bri* and *Can*) (**Fig 1A**, **Table S3**). *Ava*, *Bea* and *Caia* have a similar structure where two identical LTR sequences flank an internal domain. The internal domain contains two ORFs: one encoding for a putative POL and one encoding for an *S. stipitis*-specific protein that we named LTR-Associated Protein (Lap1 in *Ava*, Lap2 in *Bea* and Lap3 in *Caia*). Homology search failed to identify any GAG gene associated with the *Ava*, *Bea* and *Caia* retrotransposons. As Gag proteins are poorly conserved among different organisms, we hypothesise that the Lap proteins are Gag proteins.

*Ace*, *Bri* and *Can* are LINE elements composed of the Non-Coding regions NC-1 and NC-2 surrounding an internal coding region encoding for a Pol enzyme and an *S. stipitis*-specific LINE Associated protein (Linea1 in *Ace*, Linea2 in *Bri* and Linea3 in *Can*). Linea1 and Linea2, but not Linea 3, have a Zinc-Finger DNA binding motif (**Table S3**). Comparison across the CTG (Ser1)-clade revealed that *S. stipits* TE repertoire is typical of this clade. Indeed, retrotransposons are common in this yeast group: the genome of all species analysed contains LTR elements, whereas LINE elements are present in 6/8 species (**Fig 1B, Table S4**). Similarly to other CTG-(Ser1) clade yeast, we did not detect any DNA transposons integrated into the *S. stipitis* genome (**Fig 1B, Table S4**).

Our repeat analysis demonstrates that the terminal sequences of *S. stipitis* chromosomes are repeat-rich and composed of two elements with different degree of repetitiveness: telomere proximal-repeats and subtelomeric regions. The telomeric repeats are non-canonical and composed of 24-nucleotide units repeated in tandem. Each unit contains a TG motif reminiscent of typical telomeric repeats (**Fig 1C**). *S. stipitis* subtelomeric regions (the ~30KB region adjacent to telomeric repeats) are enriched in retrotransposon-derived elements. Indeed, DNA sequences with homology to *Bea* LTR-retrotransposons and *Ace* LINE-elements are found in 5/16 subtelomeric regions (**Fig 1D**, **Table S3**). No full length-retrotransposons are detected at these genomic locations. Subtelomeric regions contain several gene families members, including gene encoding for ATP-dependent DNA helicases (found in 7/16 subtelomeres), fungal-specific transcription factors (8/16 subtelomeres), MFS transporters (8/16 subtelomeres) and Agglutinine-like proteins (11/16 subtelomeres) (**Fig 1D, Table S5**) (39). In summary, our analysis demonstrates that the *S. stipitis* genome contains several classes of repetitive elements that could be major contributors of genome plasticity.

### *S. stipitis* natural isolates have distinct genomic organisations

Having identified *S. stipitis* DNA repeats, our next step was to examine *S. stipitis* phenotypic and genotypic diversity across a geographically diverse set of strains (n=27) that were collected in different habitats (**Table S1** source NRRL and NCYC collection), and that includes the sequenced Y-11545 strain (31). rDNA fingerprinting confirm that all isolates belong to the *S. stipitis* species (D1/D2 domain of the S26S rDNA similarity >99 %) (**Table S6**). Phenotypic analyses established that the natural isolates vary in their ability to utilise and grow on different carbon sources. Indeed, when compared to the reference Y-11545 strain, different natural isolates cultured in Synthetic Complete media containing the hexose sugar Glucose (SC-G), the pentose sugar Xylose (SC-X) or a mixture of both sugars as found in lignocellulose (SC-G+X) display distinct growth rate, maximum culture density and lag phase (**Fig 2A** and **Table S7**). To determine whether the natural isolates have distinct genomic organisations, we analysed their karyotype by chromosomes Contour-clamped Homogenous Electric Field (CHEF) gel electrophoresis, a technique allowing chromosome separation according to size. The CHEF electrophoresis analysis reveals clear differences in chromosome patterns demonstrating that *S. stipitis* natural isolates have a genome organised in different-sized chromosomes (**Figure 2B**). We concluded that intra-species phenotypic and genotypic variation is a common feature of *S. stipitis*.

**Figure 2.**
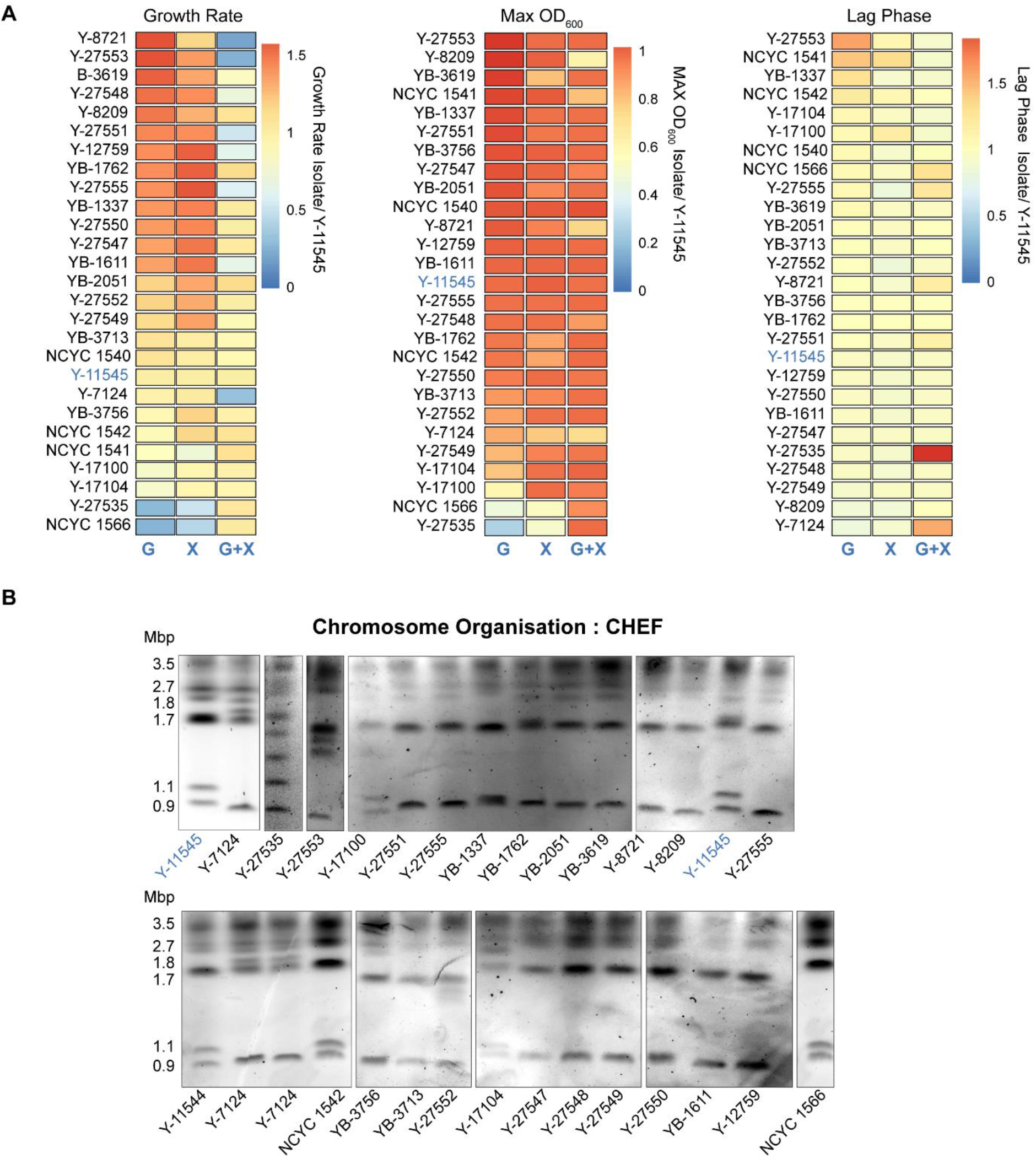
Phenotypic and Genotypic Diversity in *S. stipitis* **A)** Heatmaps comparing growth rate (Left), maximum OD (Middle) and lag time (Right) for each *S. stipitis* natural isolate in comparison to the reference Y-11545 strain (blue). Analyses were performed in Glucose (G), Xylose (X) and Glucose/Xylose (G+X) media. The heatmap data are the average of 3 biological replicates. **B)** Karyotyping of *S. stipits* natural isolates by CHEF electrophoresis. Y-11545 strain is highlighted in blue and the size of its eight chromosome is indicated.

### Hybrid genomic sequencing identifies transposable elements as drivers of *S. stipitis* genome plasticity

To date, only one *S. stipitis* isolate (Y-11545) has been sequenced and assembled at chromosome level (31). To gain insights into *S. stipitis* genetic diversity, we generated a chromosome-level sequence assembly of a second *S. stipitis* natural isolate (Y-7124) by combining MinION Nanopore with Illumina genome sequencing. This isolate was chosen because *(i)* karyotypic analysis reveals that its genomic organisation is distinct from the genomic organisation of the reference strain Y-11545, and *(ii*) Y-7124 is widely used both for industrial applications and for basic research (61).

The Y-7124 genome was sequenced to 186.88x coverage resulting in a 15.69 Mb assembly arranged in 10 contigs (**Table S8**). High accuracy reads from Illumina-sequencing enabled the correction of errors that are associated with the MinION technology. A final chromosome-level assembly was produced by manually identifying overlapping regions between contigs. Comparing the Y-7124 and Y-11545 nucleotide sequences reveals that the two natural isolates overall share a similar coding DNA sequence. The total number of SNPs between the two natural isolates is 50,495 SNPs, equating to one variant every 306 bases. The majority of these SNPs are synonymous changes (16,294 =74.25%), while ~25% (5,622) of SNPs are missense and only (0.13% (28) are nonsense (**Table S9**). Despite this high DNA sequence similarity, the Y-7124 genome is organised in eight chromosomes with different sizes and organisations from that of Y-11545 (**Fig 3A**).

**Figure 3.**
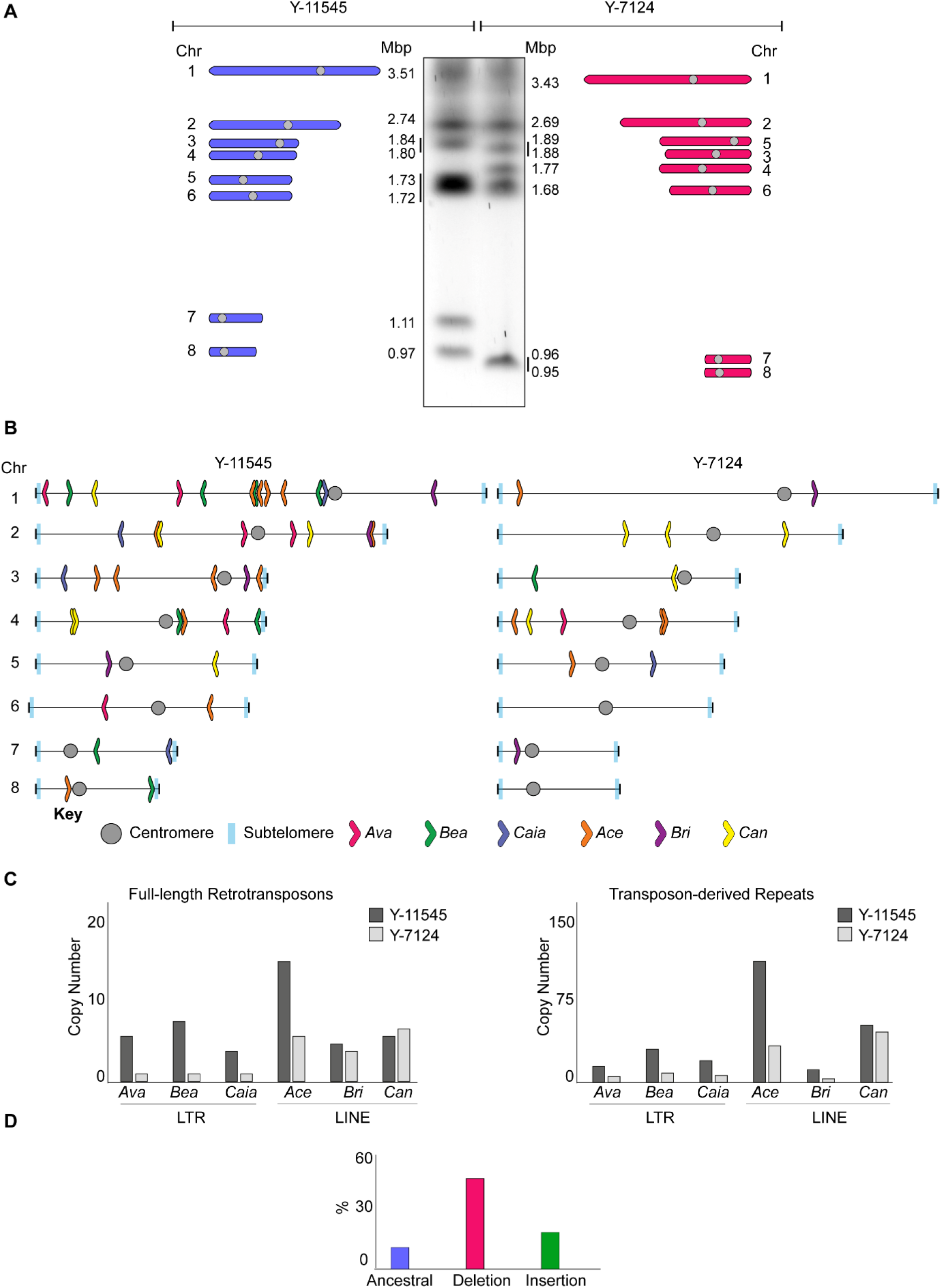
Differences in TE distribution and organisation **A)** The genomic organisation of Y-11545 and Y-7124 is distinct. *Left:* Schematics of Y-11545 chromosome organisation. Chromosome (Chr) number and size (Mbp) is indicated. *Middle:* Karyotyping of *S. stipits* Y-11545 and Y-7124 strains by CHEF electrophoresis. *Right:* Schematics of Y-7124 chromosome organisation. Chromosome (Chr) number and size (Mbp) is indicated **B)** Schematics of non-centromeric transposon family distribution in Y-11545 (*left*) and Y-7124 (*right*) **C)** Copy Number of full-length transposons (Left) and transposon-associated repeats associated with the Y-11545 (dark grey) and Y-7124 (light grey) genome. **D)** Percentage (%) of Ancestral, Deletion and Insertion sites associated with the Y-11545 and Y-7124 genomes

Comparison of the Y-7124 and Y-11545 genomes establishes that retrotransposons are significant drivers of *S. stipitis* genome diversity as one of the most prominent differences between the two genomes is the abundance and localisation of these retrotransposons (**Fig 3B**). Indeed, the number of LTR and LINE non-centromeric retrotransposons and transposons-derived repeats is greater in the Y-11545 reference genome compared to the Y-7124 genome: retrotransposons, solo LTR and truncated LINE elements account for approximately 2% of the reference Y-11545 genome and only for ~1% of the Y-7124 genome (**Fig 3C**). We classified retrotransposons loci present in both isolates (ancestral loci), those present in the reference Y-11545 genome but absent in Y-7124 (deletion loci) and those not present in the reference genome but present in a given strain (insertion loci). Out of 69 transposons loci, only ten ancestral loci (~15%) were detected in the two isolates. These sites are likely to be inactive transposons or transposons that rarely transpose. In addition, we detected 42 deletion loci (60 %) and 17 (24%) insertion loci (**Fig 3D**). The presence of deletion and insertion loci suggests that *S. stipitis* LTR transposons and LINE elements are active and competent of transposition. Although active transposons can insert into genes to cause functional consequences (62), we did not detect any TE-driven alteration in coding regions.

### Transposable Elements are sites of chromosome rearrangements

Comparison of the Y-11545 and Y-7124 genome reveals that transposon-rich regions are sites of complex chromosome rearrangements. Indeed, a transposon-rich region is the breakpoint of a reciprocal translocation between chromosome 5 and chromosome 7. This translocation causes the size change of chromosome 5 ^Y-7124^ and chromosome 7 ^Y-7124^ detected by CHEF karyotyping (**Fig 4A**). Southern analyses with a probe specific for chromosome 5 ^Y-11545^ confirms this finding (**Fig S1**). The evolutionary history of Y-11545 and Y-7124 is unknown, and therefore it is difficult to predict the molecular events underlying these genomic changes. However, sequence analysis of the rearrangement breakpoint reveals that this structural variation occurs in a genomic region that *(i)* contains homologous sequences between chromosome 5 and 7 and *(ii)* is transposon-rich and contains two inverted repeats on chromosome 7 (**Fig 4B**). A second significant difference between the genome organisation of Y-11545 and Y-7124 is found at subtelomeric regions: these regions differ in the number and organisation of subtelomeric gene families and in the number of transposon-associated repeats (**Fig 4C**). Lastly, we detected a distinct centromeres organisation where the numbers of *Tps5* retrotransposons, LTRs and LARD regions differ between the two isolates (**Fig 4D**). The presence of transposons and transposon-derived repeats associated with all these genomic locations strongly suggest that retrotransposons have mediated the chromosomal rearrangement by recombination-mediated mechanisms. Therefore, changes in transposons organisation are responsible for the bulk of genomic changes identified in two different natural isolates.

**Figure 4.**
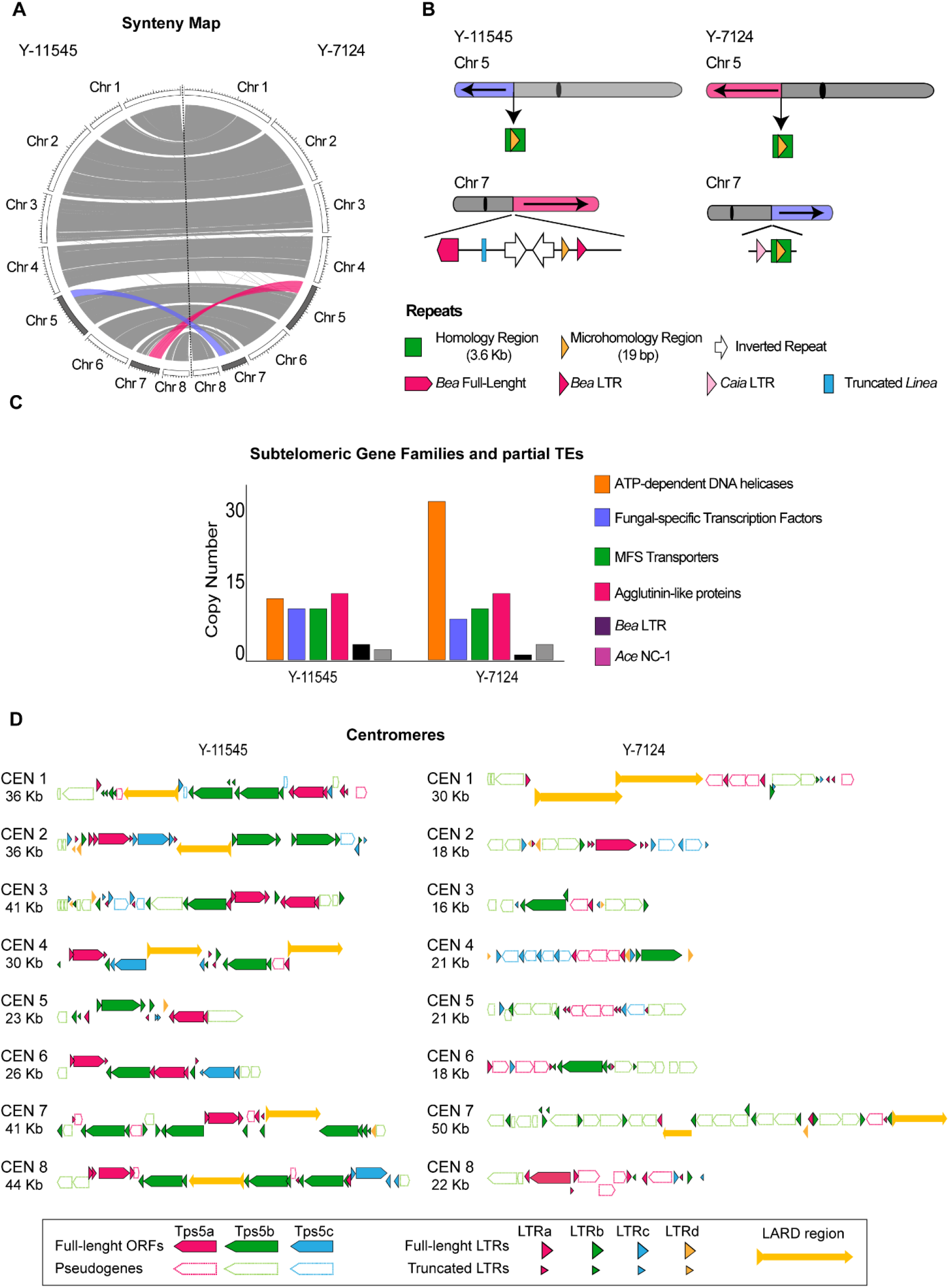
Chromosome rearrangements between *S. stipitis* natural isolates **A)** Circos plot displaying macrosynteny between Y-11545 (Left) and Y-7124 (Right). Chromosome (Chr) number and size is indicated. Recriprocal translocation between the two genomes is highlighted in purple and pink. **B)** Schematics of repetitive sequences associated with the translocation junction in the Y-11545 (Left) and Y-7124 (Right) genomes. **C)** Subtelomeric gene families and TEs distribution in the Y-11545 and Y-7124 genomes **D)** Schematics of centromere organisation in the Y-11545 (Left) and Y-7124 (Right)

### *S. stipitis* real-time evolution leads to extensive genomic changes

Our results demonstrate that intraspecies genetic diversity is common in *S. stipitis*. However, as the evolutionary history of the analysed natural isolates are unknown, it is difficult to predict whether the observed genomic changes are due to the selection of rare genomic rearrangements events. To determine the time scale of *S. stipitis* genome evolution, we investigated the genome organisation of 72 single colonies passaged daily for 8 weeks (56 passages, ~672 divisions) in SC-G+X, as its sugar composition resembles what found in lignocellulose (29) (**Fig 5A**). Strains were grown at 30 °C, a temperature that does not lead to any growth defect, and 37 °C, a stressful temperature that strongly inhibits *S. stipits* growth (**Fig 5B**). CHEF gel electrophoresis was conducted to identify possible changes in the chromosome organisation of the evolved strains. This analysis identifies genome rearrangements in 19/36 strains evolved at 30 °C and 12/36 strains evolved at 37 °C (Blue and Magenta- **Fig 5C**). Thus, changes in chromosome organisation were detected in the presence (37 °C) or absence (30 °C) of stress. To test whether chromosome rearrangements are associated with a fitness benefit, we tested the ability of the parental and 37 °C-evolved strains to grow in SC-G+X media at permissive (30 C) and restrictive (37 C) temperature (**Fig 5D**). This analysis demonstrates that 37 °C-evolved strains with no chromosomal rearrangement grow poorly at 37 °C (**Fig 5D**). In contrast, 5/12 37 °C-evolved strains with chromosome rearrangements grow better than the parental strain at this restrictive temperature (**Fig 5D**). This result suggests that changes in chromosome organisation have an adaptive value. Thus, genome plasticity is a defining feature of the *S. stipitis* genome, and its genome can rapidly change in mitotic cells propagated *in vitro*. Our results strongly suggest that the extensive genomic changes can lead to adaptation to hostile environments.

**Figure 5.**
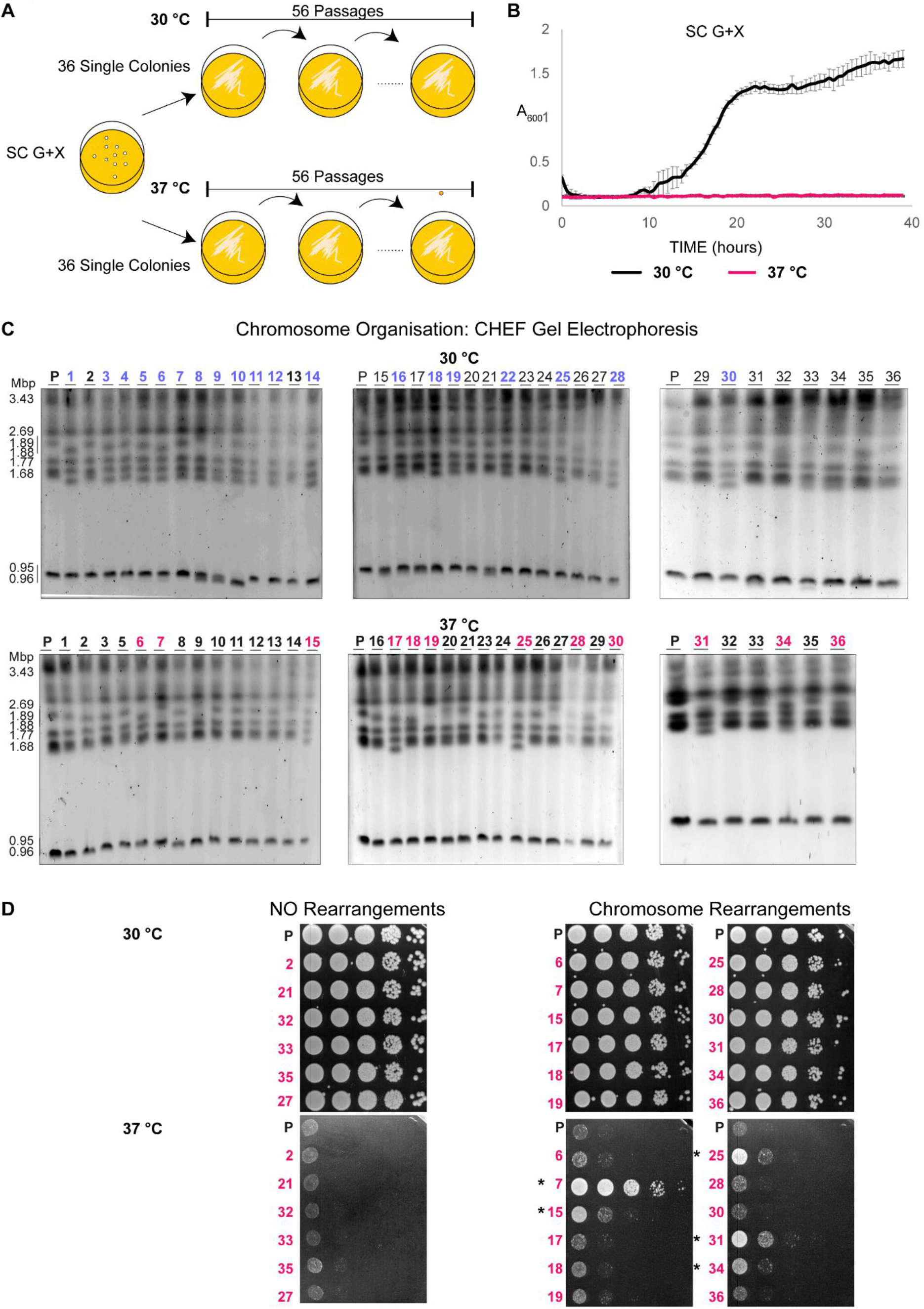
The *S. stipitis* genome is plastic following real-time evolution **A)** Schematics of laboratory evolution strategy **B)** S. stipitis growth curve in SC G+X liquid media at permissive (30 °C) and restrictive (37 °C) temperature **C)** Karyotype organisation of *S. stipitis* colonies following 8 weeks of laboratory evolution at 30 °C and 37 °C. Variation in the structure following laboratory evolution at 30 °C (blue) and 37 °C (magenta) is indicated **D)** Serial dilution assay showing growth of parental (P) and 37 °C-evolved strains without (NO Rearrangements) and with (Rearrangements) at 30 °C and 37 °C. The CHEF analysis strain number is indicated (Magenta). * indicates colonies with a fitness advantage compared to the parental strain.

## DISCUSSION

Here we demonstrate that the yeast *S. stipitis* has a plastic genome and that genome plasticity is linked to adaptation to hostile environments. We show that non-centromeric retrotransposons are significant drivers of *S. stipitis* genome diversity. These findings have important implications for developing economically viable second-generation biofuels and better understanding the CTG (Ser1)-clade biology.

### Retrotransposon are drivers of *S. stipitis* genome diversity

Our repetitive sequence analysis demonstrates that *S. stipitis* has a DNA repeats content typical of the CTG (Ser1)-clade including TEs, non-canonical terminal telomeric repeats and subtlomeric regions. As observed in other members of the CTG (Ser1)-clade (60), we did not detect any DNA-transposons or MRS repeats. One of our major findings is that that non-centromeric retrotransposons are significant drivers of *S. stipitis* genome diversity. Our data support the hypothesis that *S. stipitis* TEs generate genome diversity via two distinct mechanisms: transposition into new genomic locations and recombination-mediated chromosome rearrangements. Indeed, we demonstrated that the number and genomic position of non-centromeric retrotransposons vary between the Y-11545 and Y-7124 *S. stipitis* isolates. Significantly, we did not detect transposon insertions into coding regions. However, transposons might alter *S. stipitis* gene expression by inserting into gene regulatory regions (62). We propose that *S. stipitis* transposons are active and generate genome diversity by jumping into different genomic locations. Our data also indicate that TEs can generate further genome diversity though either homologous recombination of nearly identical TE copies or by faulty repair of double-strand breaks generated during transposable elements excision (62). Indeed, we find that the translocation breakpoint between chromosome 5 and chromosome 7 is enriched in retrotransposons. Furthermore, TE-rich subtelomeric regions and centromeres have a distinct organisation in the two analysed isolates suggesting that the transposons drive this genetic diversity. We hypothesise that transposons elements cause the genetic variability observed during laboratory passaging. In the future, it will be important to apply the hybrid genome sequencing approaches presented in this study to dissect the nature of these rearrangements.

### Genome plasticity and production of second-generation biofuels

One of our key findings is that the *S. stipitis* genome is intrinsically plastic and that chromosome rearrangements are frequent events under stress or unstressed conditions. Second-generation biofuels, generated by fermentation of agriculture and forestry waste, have an enormous potential to meet future energy demands and significantly reduce petroleum consumption. To meet the requirements for industrial applications, second-generation biofuels need to be generated by microorganisms that can efficiently utilise and ferment all the sugars found in lignobiomass (63). Consequently, *S. stipitis* is one of the most promising yeast for producing second-generation bioethanol as it can efficiently ferment both hexose and pentose sugars (25, 26, 29). However, robust economically viable *S. stipitis* platforms still require significant development as this organism struggles to survive under the harsh environments generated during second-generation biofuel production. For example, *S. stipitis* growth and fermentation is inhibited by the chemical pre-treatment required to extract glucose and xylose from lignobiomass (61). Growth is also inhibited at high ethanol concentration, and *S. stipitis* ferments xylose less efficiently than glucose. Evolutionary engineering approaches under selective conditions (i.e. presence of inhibitory compounds, high concentration of xylose or ethanol) have been applied to isolate better performing *S. stipitis* strains (61).

Our data predict that the genetic make-up and associated improved phenotypes of superior biofuel-producer strains are unstable and that the genetic drivers of improved phenotypes might be lost over time. This hypothesis could explain why short-read Illumina genome sequencing has failed to identify point mutations or indels that could explain the superior performance of *S. stipitis* strains (64). It is also possible that *S. stipitis* superior strains carry stable complex chromosomal rearrangements with a breakpoint at DNA repeats. Such rearrangements could not have been identified by Illumina sequencing as short sequenced fragments will not resolve changes associated with long repetitive elements. Thus, economically viable use of *S. stipitis* for second-generation biofuels production will require an in-depth analysis of the genomic structures of superior strains.

### Genome plasticity in the CTG (Ser1)-clade

The CTG-Ser1 clade is an incredibly diverse yeast group that includes many important human pathogens and non-pathogenic species (17). Our data support the hypothesis that genome plasticity is a general feature of the CTG (Ser1) yeast clade as it has been observed in *C. albicans* and *S. stipitis* ((21, 22, 65) and this study), two organisms with a very different lifestyle. Indeed, while *C. albicans* is a diploid opportunistic human fungal pathogen that lives almost exclusively in the human host, *S. stipitis* is a haploid non-pathogenic yeast found in the gut of wood-ingesting beetles hardwood forests or areas high in agricultural waste (29, 66). Furthermore, while *C. albicans* lacks a canonical sexual cycle and its associated meiosis, *S. stipitis* has a canonical sexual cycle whereby mating of haploid cells generate diploid cells that undergo meiosis and produce haploid spores (30).

Our results highlight that stress regulates genome plasticity differently in *C. albicans* and *S. stipitis*. It has been demonstrated that stress exacerbates *C. albicans* genome instability (21, 67). In contrast, we found that *S. stipitis* genome instability is not regulated by stress as we detected a similar rate of chromosomal rearrangements when cells are continuously passaged in unstress (30 °C) or stress (37 °C) conditions. Importantly, we also demonstrated that the large genomic changes are associated with fitness benefits suggesting that genome plasticity is instrumental for adaptation to hostile environments.

In summary, our study demonstrates for the first time that *S. stipitis* genome is plastic. Understanding the cause and effect of this extensive genome plasticity is of paramount importance to understand the biology of the CTG(Ser1)-clade of fungi.

## DATA AVAILABILITY

This Whole Genome Shotgun project has been deposited at DDBJ/ENA/GenBank under the accession JADGGA000000000. The version described in this paper is version JADGGA010000000. Illumina and nanopore sequence data associated with this work have been deposited on the Sequence Read Archive (SRA) under BioProject PRJNA609885.

## ACKNOWLEDGEMENTS

We thank Dr Patricia Slininger, members of the Buscaino Lab, the Kent Fungal Group and Dr Jan Soetaert, for discussion and critical reading of the manuscript.

---

**Figure S1.** Southern Blot analysis confirm the Chr5 / Chr7 translocation

Left: Schematics of chromosome 5 (Chr 5) and chromosome 7 (Chr 7) in Y-11545 (Left) and Y-7124 (Right). Reciprocal translocation is highlighted in purple and pink. Southern Probe is indicated. Right: Southern Blot of Y-11545 and Y-7124 chromosomes separated by CHEF gel electrophoresis. Full chromosome profiling (EtBr) and Southern Blot results (Southern) are indicated

