## Supplementary Data for "The genome of the CTG(Ser1) yeast S*cheffersomyces stipitis* is plastic" (from another version of this preprint: version 1, 2021-02-16)

**Supplementary Table S1**: List of Strains used in this study

| **Collection** | **Identifier** | **Habitat** | **Region** | **Country** |
| --- | --- | --- | --- | --- |
| NRRL | Y-11545 (1) | Unknown | Delft | The Netherlands |
| NRRL | Y-7124 | Insect (IN) larva on fruit tree | Rhône, Lyon | France |
| NCYC | 1540 | Habitat Insect larva | Rhône, Lyon | France |
| NCY | 1542 | Insect (IN) larva on fruit tree | Rhône, Lyon | France |
| NCY | 1566 | Insect (IN) larva on fruit tree | Rhône, Lyon | France |
| NRRL | Y-17104 | Industrial fermentations (IFM), from xylose fermentation | Delft | The Netherlands |
| NRRL | YB-3756 | Pine and coniferous trees (PTR), gymnosperms, dead pine tree, | Gainesville, Florida | USA |
| NRRL | Y-27547 | Insect (IN), beetle, Odontotaenius disjunctus, Passalidae, Burke Co. Shell | Georgia | USA |
| NRRL | Y-27548 | Insect (IN), beetle, Odontotaenius disjunctus, Passalidae, | Lake Herrick Park, Georgia | USA |
| NRRL | Y-27549 | Insect (IN), beetle, Passalidae, | Orangeburg County, South Carolina | USA |
| NRRL | Y-27550 | Insect (IN), beetle, Odontotaenius disjunctus, Passalidae, | Burden, Baton Rouge, Louisiana | USA |
| NRRL | YB-1611 | Hard wood trees (HTR), angiosperms, shagbark hickory, | Peoria, Illinois | USA |
| NRRL | Y-12759 | Soil or rock (SL), forest soil, | Georgia | USA |
| NRRL | YB-3713 | Frass or Insect tunnels (FR), frass on dead oak log, | Gainesville, Florida | USA |
| NRRL | Y-27552 | Insect (IN), beetle, Verres sternbergianus, Passalidae, | Barro Colorado Island | Panama |
| NRRL | Y-27535 | Insect (IN), gut of a Passalid beetle, | Kansas | USA |
| NRRL | Y-8209 | Tree, unknown type (TR), tree trunk, | New Orleans, Louisiana | USA |
| NRRL | Y-8271 | Frass or insect tunnels (FR), frass, | New Orleans, Louisiana | USA |
| NRRL | Y-17100 | Insect (IN), insect larvae on fruit tree, | Rhône, Lyon | France |
| NRRL | Y-27551 | Insect (IN), beetle, Odontotaenius disjunctus, Passalidae | Duches Dr. Park, Baton Rouge, Louisiana | USA |
| NRRL | Y-27555 | Insect (IN), beetle, Odontotaenius disjunctus, Passalidae, | Oxford, Pennsylvania | USA |
| NRRL | YB-1337 | Tree, unknown type (TR), rotted log, | Wohlwend Farm, Marion, Illinois | USA |
| NRRL | YB-2051 | Frass or insect tunnels (FR), frass, dead black oak. | Missouri | USA |
| NRRL | YB-3619 | Frass or insect tunnels (FR), frass, American elm. |  | USA |
| NRRL | Y-27553 | Insect (IN), beetle, Verres sternbergianus, Passalidae, | Barro Colorado Island | Panama |
| NRRL | YB-1762 | Frass or insect tunnels (FR), frass, rotten logs, Brownfield Woods | Illinois | USA |
| NCYC | 1541 | Insect (IN) larva on fruit tree | Rhône, Lyon | France |

**Supplementary Table S2:** Identification of S. stipitis natural isolates by Sanger sequencing (D1/D2 domain of 26S rDNA gene)

| **Code** | **Blast result:**  **D1/D2 domain**  **of 26S rDNA gene** | **% identity** | **E-value** |
| --- | --- | --- | --- |
| ATCC 58784 | *Scheffersomyces stipitis* | 99.47 | 0.0 |
| ATCC 58376 | *S. stipitis* | 99.14 | 0.0 |
| ATCC 62970 | *S. stipitis* | 98.63 | 0.0 |
| ATCC 62971 | *S. stipitis* | 99.65 | 0.0 |
| NRRL Y-17104 | *S. stipitis* | 99.65 | 0.0 |
| NRRL YB-3756 | *S. stipitis* | 99.65 | 0.0 |
| NRRL Y-27547 | *S. stipitis* | 99.82 | 0.0 |
| NRRL Y-27548 | *S. stipitis* | 99.65 | 0.0 |
| NRRL Y-27549 | *S. stipitis* | 99.47 | 0.0 |
| NRRL Y-27550 | *S. stipitis* | 99.82 | 0.0 |
| NRRL YB-1611 | *S. stipitis* | 99.82 | 0.0 |
| NRRL Y-12759 | *S. stipitis* | 99.65 | 0.0 |
| NRRL YB-3713 | *S. stipitis* | 99.13 | 0.0 |
| NRRL Y-27552 | *S. stipitis* | 99.65 | 0.0 |
| NRRL Y-27535 | *S. stipitis* | 99.65 | 0.0 |
| NRRL Y-8209 | *S. stipitis* | 99.82 | 0.0 |
| NRRL Y-8271 | *S. stipitis* | 99.47 | 0.0 |
| NRRL Y-11545 | *S. stipitis* | 99.48 | 0.0 |
| NRRL Y-17100 | *S. stipitis* | 99.65 | 0.0 |
| NRRL Y-27551 | *S. stipitis* | 99.47 | 0.0 |
| NRRL Y-27555 | *S. stipitis* | 99.29 | 0.0 |
| NRRL YB-1337 | *S. stipitis* | 99.82 | 0.0 |
| NRRL YB-1762 | *S. stipitis* | 99.65 | 0.0 |
| NRRL YB-2051 | *S. stipitis* | 99.82 | 0.0 |
| NRRL YB-3619 | *S. stipitis* | 99.31 | 0.0 |
| NRRL Y-7124 | *S. stipitis* | 99.65 | 0.0 |
| NRRL Y-27553 | *S. stipitis* | 99.82 | 0.0 |

**Supplementary Table S3.** Quality assessment of Y-7124 Genome sequencing

| N. of contigs | 11 |
| --- | --- |
| Total length (Mbp) | 15.69 |
| Largest contig (Mbp) | 2.69 |
| N50 (Mbp) | 1.88 |
| N75 (Mbp) | 1.67 |
| L50 | 4 |
| L75 | 6 |

Notes

**N50:** Total length of the shortest contig at 50% of the genome length (50% of the genome is contained in contigs larger than this value).

**L50:** Smallest number of contigs whose length sum up makes half of the genome size.

**Supplementary Table S4:** Primers used in this study

| AB798 | GCATATCAATAAGCGGAGGAAAAG |
| --- | --- |
| AB799 | GGTCCGTGTTTCAAGACGG |
| AB1028 | CAACTTCAAACACCGGCTCG |
| AB1029 | CTGGTGTCGACGGTAAACCA |
